# Inadequate level of knowledge, mixed outlook and poor adherence to COVID-19 prevention guideline among Ethiopians

**DOI:** 10.1101/2020.07.22.215590

**Authors:** Edessa Negera, Tesfaye Moti Demissie, Ketema Tafess

**Affiliations:** Hawassa University, Department of Biology, Ethiopia; College of Health Sciences, Arsi University, Ethiopia; Medical Science Division, Centre for Vaccinology and Tropical Medicine, University of Oxford

## Abstract

COVID-19 has a potential to cause chaos in Ethiopia due to the country’s already daunting economic and social challenges. Living and working conditions are highly conducive for transmission, as people live in crowded inter-generational households that often lack running water and other basic sanitary facilities. Thus, the aim of this study was to investigate the knowledge, attitudes and practices (KAP) of Ethiopians toward COVID-19 following the introduction of state of emergency by the Ethiopian government to curb the spread of the disease. A cross-sectional study design was conducted in nine reginal states and two chartered cities. Data for demographic, Knowledge, attitude and practice toward COVID-19 were collected through telephone interview from 1570 participants. Descriptive and bivariate analyses using chi-square test, t-test or analysis of variance were performed as appropriate. Binary and multiple logistic regression analysis were used to measure the relationship between the categorical dependent variables and one or more socio-demographic independent variables with two-tailed at α=0.05 significance level and 95% of confidence interval. The level of good knowledge, favourable attitude and good practice among the respondents were 42%, 53.8% and 24.3% respectively. Being rural resident, older than 50 years, having at least primary education, being resident of Amhara and Oromia regions were independent predictors of knowledge level. While being rural resident, married, employed, having at least basic education, being residents of Afar, Amhara, Gambela, Oromia and Somali regions were found to be the best predictors of the attitude, being rural resident, government employee, having at least basic education, and living outside of the capital were the independent predictors of practice level of the respondents. The finding revealed that Ethiopians have inadequate level of knowledge and are generally have a mixed outlook on overcoming the pandemic with poor adherence to COVID-19 prevention practice. reinforcing preventive measures and intensifying sensitization campaigns to fill the knowledge gap and persuading people to follow the preventive measures set by the government with concurrent evaluation of the impacts of these measures on knowledge and practice is highly recommended to mitigate the disease.

## 1. Introduction

Coronavirus disease 2019 (COVID-19) is a pandemic disease caused by a novel human coronavirus (SARS-COV-2) first identified in china in December 2019 (1). The World Health Organization declared the outbreak of COVID-19 as a pandemic on the 11^th^ of March 2020, after it has spread to 113 countries worldwide (2).

According to current evidence, the COVID-19 virus is primarily transmitted rapidly between people through respiratory droplets and contact routes (3). Airborne transmission has been suggested by some studies (4, 5). Recent experimental studies have examined the stability of SARS-CoV-2, showing that the virus remains infectious in aerosols for hours (5) and on surfaces up to days (5, 6). The mean incubation period of COVID-19 is about 3–9 days with a range between 0–24 days (3, 7). However, the mean time between successive cases in a chain of transmission is about 3-8 days suggesting that one becomes contagious before symptoms present about 2.5 days earlier from the onset of symptoms (8). Studies estimated that about 44% of transmission of COVID-19 occur before the onset of symptoms (9).

The first case of COVID-19 was confirmed in Ethiopia on 13^th^ of March 2020 (10). The state of emergency was declared by the government on 8^th^ of April 2020 to control the pandemic. The state of emergency includes closing schools, banning public gatherings and requiring employees to work from home (10). The introduction of the state of emergency has been welcomed by most citizens and institutions but was not without critiques from some political opposition parties. While the federal and regional governments announced measures such as suspending large gatherings and inter-city public transport, authorities have not introduced a comprehensive lockdown to try to contain the virus due to some real challenges. Firstly, most citizens live day-to-day and they may see a complete lockdown as counterproductive and unfair to those on the bottom rungs of society. Secondly, complete lockdown could worsen the life of the vulnerable segments of society such as street children, internally displaced persons and refugees (11).

COVID-19 has the potential to cause chaos in Ethiopia due to the country’s already daunting economic and social challenges. On one hand, the public health risks presented by COVID-19 are vast. Living and working conditions are highly conducive for transmission, as people live in crowded inter-generational households that often lack running water and other basic sanitary facilities. Allowing economic activity to continue unchecked could lead to huge infections within months, with serious cases quickly overwhelming the already weak health system (12).

Public health intervention measures are rapidly changing around the world to cope up with the rapid transmission of the disease and to minimize the risk for infection and speared of the disease. However, miscommunication regarding the threat of COVID-19 may lead to public confusion and inaction (13). Being older and having underlining health conditions, reduced ability to access and understand health information, inability to make well-informed decisions and failure to take optimal health-promoting are among the greatest risk factors for sever infection and death due to COVID-19 (13, 14). This situation is especially true when the health information itself is not timely, trusted, consistent, or actionable particularly in sub-Saharan African countries where health inequalities are worsened by lack of political commitment and good governance (15). As a result, a wide range of health disparities may exist by age, ethnicity, regions, political affiliations and socio-demographic status (15-17). Therefore, During COVID-19 pandemic when the understanding of critical preventive measures and ever-changing public health messages is most important, many vulnerable populations may be further marginalized by inadequate health communication, posing substantial risks to themselves and their communities.

On the other hand, the success of national preventive strategy of a pandemic is largely depends on the adherence of the public to the preventive measures established by the government. The public adherence is likely to be influenced by trust in government (18) and by public knowledge and attitude toward the pandemic (19). By evaluating the public knowledge, attitude and practice about COVID-19 pandemic, it is possible to explore attributes that influence the public in adopting practices and responsive behaviour toward COVID-19 prevention measures.

Thus, the aim of study was to investigate the knowledge, attitudes and practices (KAP) of Ethiopians toward COVID-19 following the introduction of state of emergency by the Ethiopian government to help curb the spread of COVID-19 following 55 cases and 2 fatalities on April 8, 2020 (20). we conducted a time sensitive KAP study in all regional states and chartered cities of Ethiopia from May 20, 2020 to June 20, 2020.

## 2. Materials and methods

### 2.1. Study design and site

A cross-sectional study was conducted from May 20 to June 20, 2020 in nine regional states and two charted cities of Ethiopia. The regional statues include Tigray, Afar, Amhara, Oromia, Harari, Somali, Gambela, Benishangul and Southern Nations, Nationalities and Peoples’ Region. (SNNP). The two charted cities are Addis Ababa (Finfinnee) and Dire Dawa.

### 2.2. Study population

The study population included all peoples living in the nine regional states and two chartered cities who are 18 years old and above during the study period.

### 2.3. Study procedure

This study was conducted after the government declared a state of emergency in the country to help curb the spread of COVID-19 following 55 cases and 2 fatalities on April 8, 2020. As face to face data collection was impossible, we opted to use telephone interview for enrolling potential participants. We ruled out the online enrolling methods for two reasons: Firstly, only about 17% of the population has access to the internet penetration but more than 45% of the population has access to either mobile network or fixed landline telephones in 2020 (21). Secondly, adult literacy is only 52% and social media penetration in Ethiopia is 5.5% by January 2020 (22). As a result, telephone interview was found to be the best alternative among the existing methods of data collection where face-to-face data collection is not possible.

### 2.4. Study tools

We adapted the survey tools from previous KAP studies towards COVID-19 (2, 13, 19). We included four core themes to assess the KAP of Ethiopians toward COVID-19. (i) demographic data collection tools. (ii) knowledge about COVID-19, (iii) attitude toward COVID-19 and (iv) practices appropriate to curb covid-19 infection and transmission.

#### Demographic questions

Each respondent was asked for age, gender, residence, occupation, region, educational and marriage status.

#### COVID-19 Knowledge

Demonstrated knowledge of COVID-19 was assessed through asking the respondents 10 close-ended questions. The questions include the respondents’ knowledge about the cause, transmission route, clinical symptoms, prevention and control measures of COVID-19. Respondents were given three response options: ‘true’, ‘false’ or ‘I do not now’. A correct response was assigned 1 point, while an incorrect or I do not know response to the question was assigned 0 points. Each respondent achieves between 0 and 10 score points. Higher score indicates better knowledge of COVID-19 (supplementary (1).

#### Attitude toward COVID-19

To assess the attitude of respondents toward COVID-19, we used 5 questions. Two questions include confidence of the respondent on COVID-19 prevention methods and confidence on the government to contain the spreading of coronavirus. They were also asked if they are in favour of the state of emergency introduced to curb the spread of COVID-19, their perception to get infected with coronavirus and if they believe traditional herbs and religious faith can cure COVID-19. For each question a 4-point Likert scale response options (strongly disagree, disagree, agree and straggly agree) were provided (supplementary (1).

#### Practice towards COVID-19 prevention

To investigate how aften respondents’ practice COVID-19 prevention, 5-questions were used. The questions include how often the respondent wash its hand with soap, avoid non-essential travel, keep 2-meter social distancing, avoid social gatherings and avoid touching eyes, nose, and mouth with unwashed (unsensitized) hands. For each question a 4-point Likert scale response options (never, sometimes, usually and always) were provided.

### 2.5. Data collection methods

Twenty data collectors were recruited and trained online. The data collectors were recruited from each region. All data collectors had previous experience of KAP data collection by phone. In most cases, phone numbers were randomly selected from local phone book. Following the verbal consent, each respondent was assured that the voice is not recorded, and any personal data is not required to participate in the study.

### 2.6. Operational definitions

#### Independent variables

all sociodemographic variables: age, sex, education, marital status, residence, employment and region are considered as independent variables.

#### Dependent Variables

knowledge, attitude and practice toward COVID-19 were considered as dependent variables for all statistical analysis.

#### Good level of knowledge

Respondents who scored above the mean score of the total respondents were considered as having good level of COVID-19 knowledge and those who scored below the mean as having poor level of COVID-19 Knowledge.

#### Favourable attitude

A response of agree or strongly agree to attitude questions were considered as favourable attitude for logistic regression analysis. However, for two questions: “I believe that traditional herbs and faith such as holy water can cure COVID-19; and I do not think I will get sick from COVID-19” a response of strongly disagree or disagree were considered as favourable attitude.

#### Good practice

a response of “usually” and “always” to each practice question was considered as having good practice towards COVID-19 prevention.

### 2.7. Statistical analysis

Descriptive statistics (means with standard deviations and percentage frequencies) were calculated for all survey responses. Associations between socio-demographic characterises and responses to COVID-19 knowledge, attitude and practice were investigated in bivariate analyses using chi-square test, t-test or analysis of variance as appropriate. Binary and multiple logistic regression analyses were used to measure the relationship between the categorical dependent variables (knowledge, attitude and practice) and one or more socio-demographic independent variables. results of logistic regression have been reported as adjusted odds ratio (AOR). Data were analysed using Stata for Windows, version 16 (Stata Corp., College Station, TX, USA with two-tailed at α=0.05 significance level and 95% of confidence interval

## 3. Result

### 3.1. Social and demographic characteristics

A total of 1570 participants completed the survey questionnaire of which 896 (57.1%) of the respondents were male. A third of the respondents were from rural residents. The age of the respondents ranges from 18 and 73 years with a mean ± standard deviation (SD) of 31.6 year ± 12.8. Among the respondents, 660 (42.0%) had college or university education while 260 (16.6%) are unable to read and write. Occupation-wise, about a third (34.9%) of them were government employees while a fifth (19.5%) of them were self-employed and one in 6 (15.9%) were farmers (Table 1).

**Table 1.** Sociodemographic characteristics of respondents (N=1570).

| Demographic variable s |  | Number | Percentage (%) |
| --- | --- | --- | --- |
| Sex | male | 896 | 57.1 |
|  | female | 674 | 42.9 |
| Residence | Urban | 1094 | 69.7 |
|  | Rural | 476 | 30.3 |
| Ager group | 18-29 | 860 | 54.8 |
|  | 30-39 | 332 | 21.1 |
|  | 40-49 | 174 | 11.1 |
|  | 50-59 | 128 | 8.2 |
|  | 60+ | 76 | 4.8 |
| Marital status | single | 662 | 42.2 |
|  | married | 894 | 56.9 |
|  | widow | 14 | 0.9 |
| Education | illiterate | 260 | 16.6 |
|  | basic | 292 | 18.6 |
|  | primary | 140 | 8.9 |
|  | secondary | 218 | 13.9 |
|  | college/university | 660 | 42.0 |
| Occupation | farmer | 250 | 15.9 |
|  | government employee | 306 | 19.5 |
|  | self-employed | 548 | 34.9 |
|  | student | 466 | 29.7 |
| Region/chartered city | Addis Ababa | 326 | 20.8 |
|  | Afar | 42 | 2.7 |
|  | Amhara | 190 | 12.1 |
|  | Benishangual Gumuz | 46 | 2.9 |
|  | Dire Dawa | 52 | 3.3 |
|  | Harari | 46 | 2.9 |
|  | Gambela | 68 | 4.3 |
|  | Oromia | 552 | 35.2 |
|  | Somali | 60 | 3.8 |
|  | SNNPRS | 90 | 5.7 |
|  | Tigray | 98 | 6.2 |
|  | Total | 1570 | 100.0 |

### 3.2. COVID-19 Knowledge

The overall correct response of the respondents to the knowledge questions about the COVID-19 varies from 14.0% to 73.6% (Table 1) with the mean knowledge score of 4.2 (SD=2.809, range 0-10) suggesting an overall of 42% (4.2/10*100) correct rate on COVID-19 knowledge test. Knowledge scores significantly differ across each question. Nearly three-fourth (73.6%) of the respondents have heard about COVID-19 but only 42.4% and 37.8% knew COVID-19 transmission and clinical manifestations, respectively. Only 14% of the respondents knew that the asymptomatic transmission of COVID-19. While about 63.8% of the respondents had correctly answered COVID-19 prevention methods such as handwashing, social distancing, avoiding crowd gatherings and using face mask. only 18.6% of them knew that people who have COVID-19 symptoms should self-isolate at least for 7 days and their household contacts for 14 days. A third of the respondents consider that only elderly people or people with chronic health conditions develop sever condition if infected with COVID-19. Nearly 80% of the respondents did not know that currently there is no effective drug to cure COVID-19 nor vaccines to prevent the infection (Table 2).

**Table 2.**
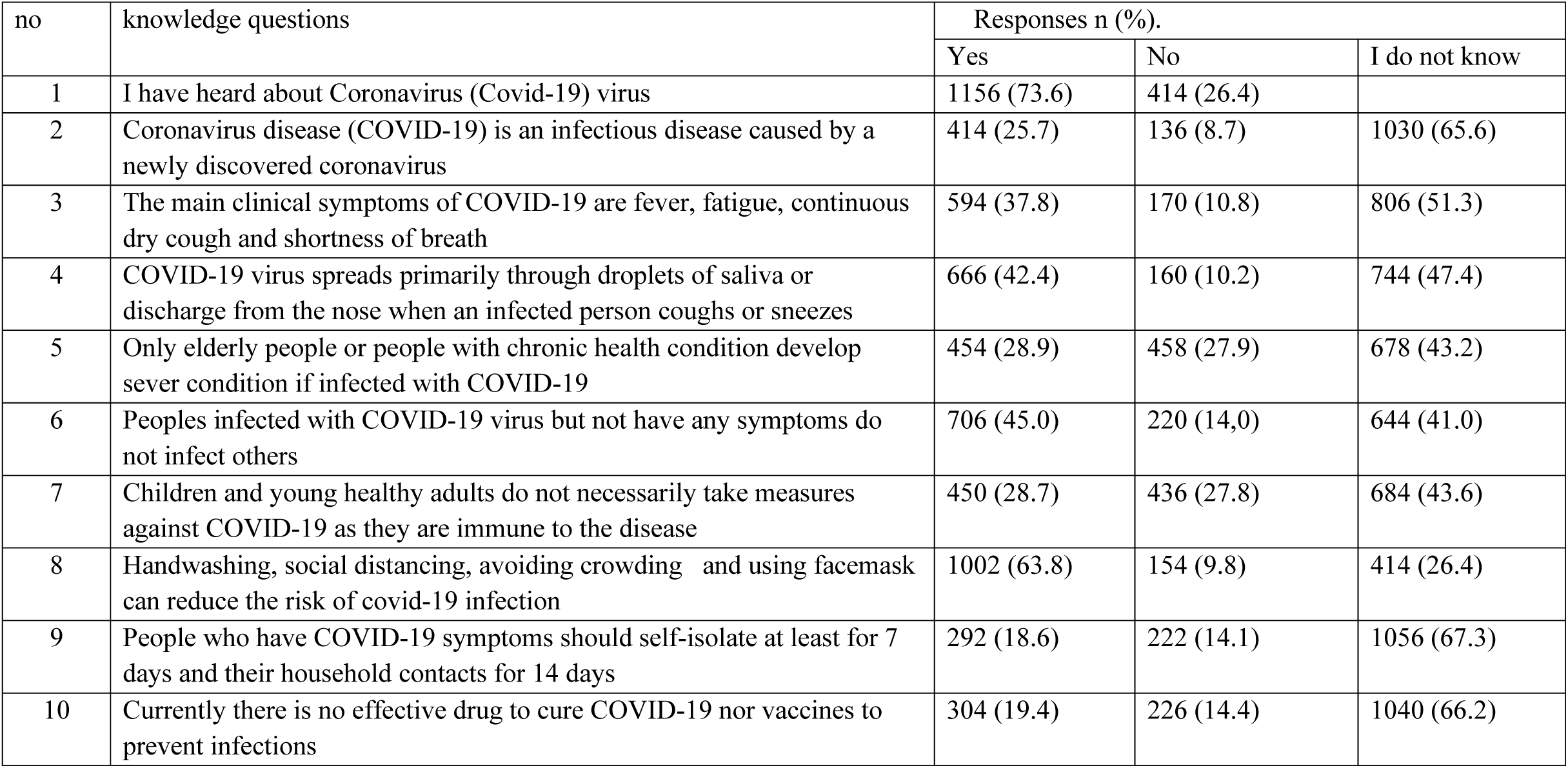
Summary of respondents’ response to COVID-19 knowledge questions (N=1570).

The student t-test has shown that the level of COVID-19 knowledge was low among female respondents compared to male respondents (P=0.05). Older respondent (≥50 years) had low COVID-19 knowledge than the other age group (F=8.38, P <0.0001). Similarly, there was a significant difference in the knowledge score of the respondents among age groups, residence, educational status, marital status and regions of residence (Table 3).

**Table 3.**
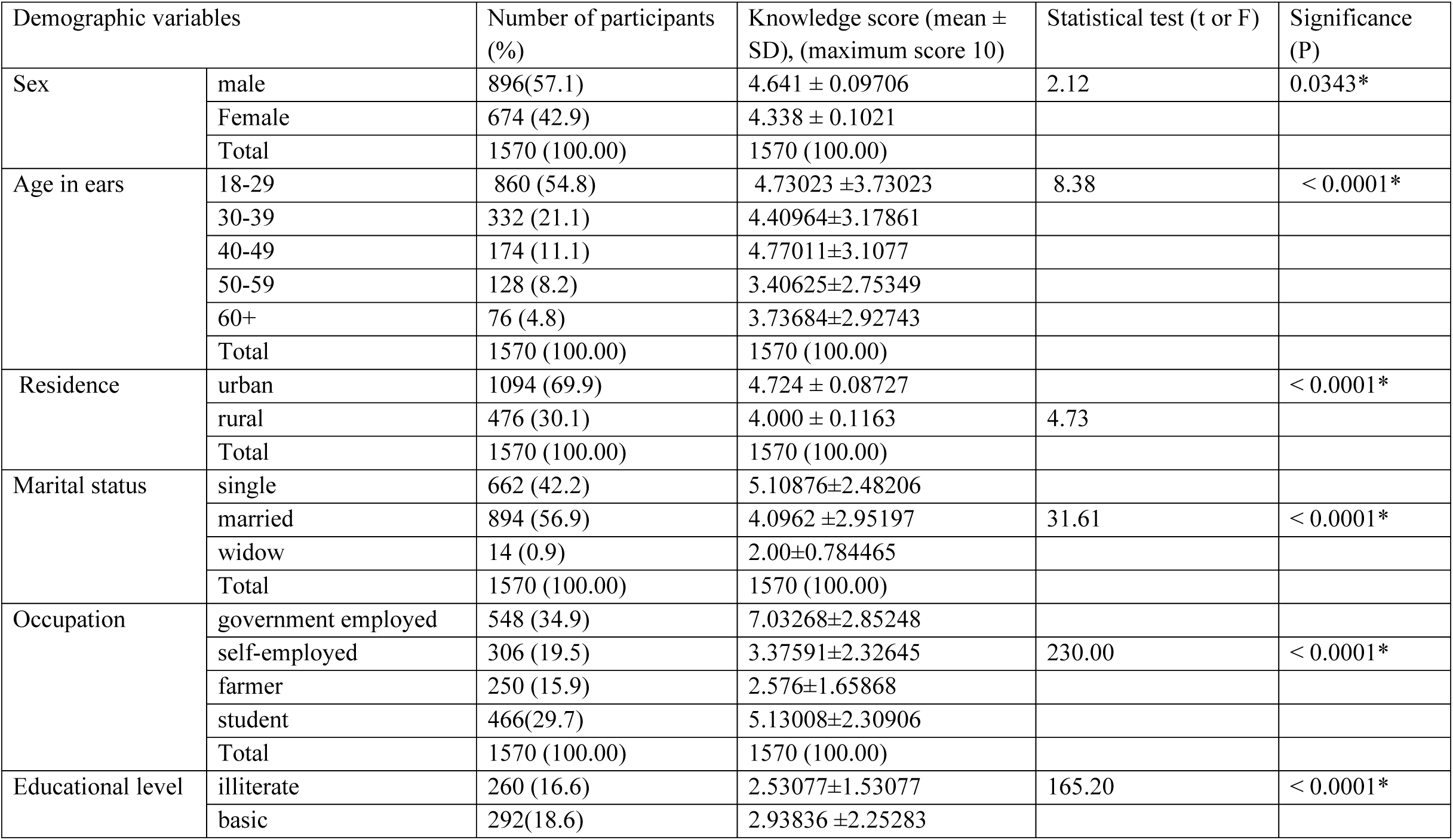
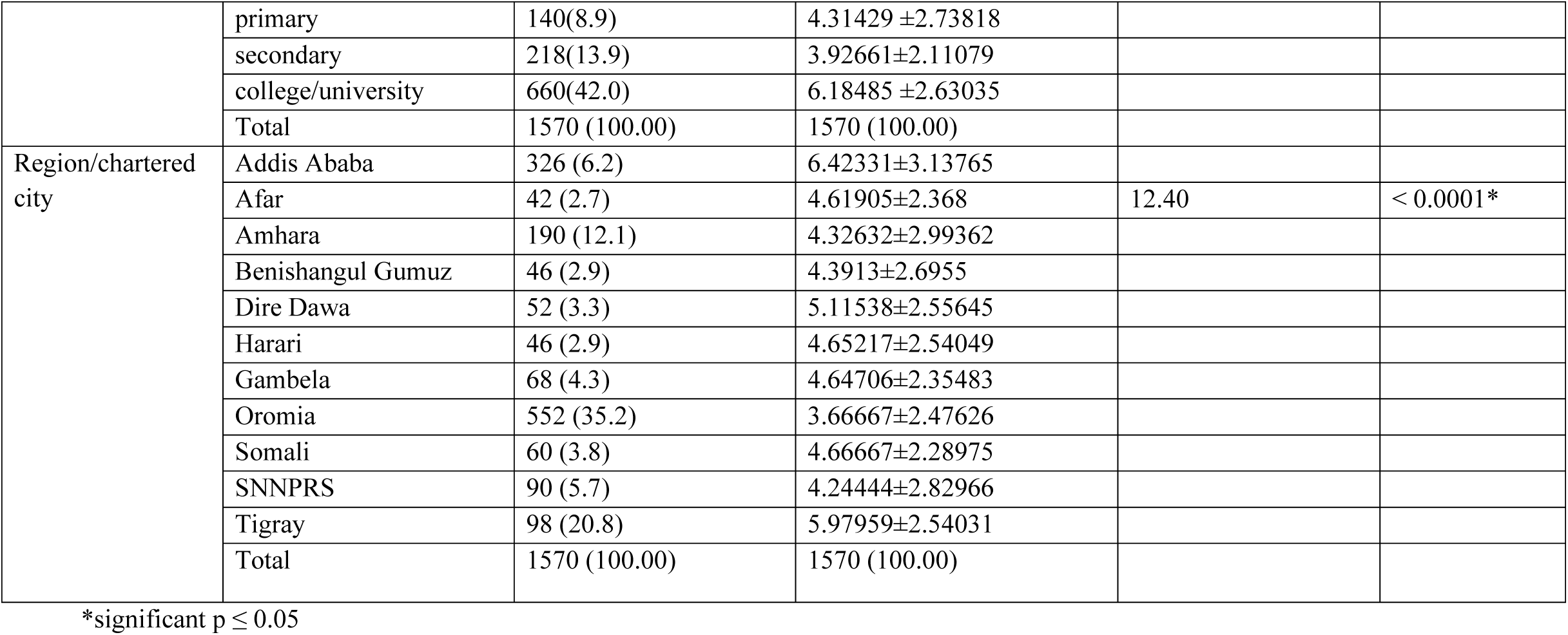
Respondents knowledge score of COVID-19 by demographic variables (N-=1750).

After grouping respondents into good (mean score ≥ 4.2) and poor (mean score < 4.2) level of knowledge using a bloom’s cut of points, binary and multivariable logistic regression was performed. Age, occupation, residence and education have been significantly associated with COVID-19 knowledge of the respondents with correct prediction of 72.2% in multivariable regression analysis. Multivariable logistic regression analysis has shown that respondents of age group 50-59 (AOR= 0.562; (95% CI: 0.4285-0.708) and above 59 years old (AOR=0.387; (95%CI: 0.1924-0.4982) were less knowledgeable about COVID-19 compared to respondents in the age group of 18-29. Similarly, respondents those who had college or university education had more than 14 times higher knowledge (AOR=14.026; (95%CI: 9.56431-21.8764) compared to those who are unable to read and write. Respondents from the two most populous regions, Amhara (AOR=0.497 (95%CI: 0.2990-0.6542) and Oromia (AOR= 0.4512; 95%CI: 0.3218-0.8026) regional states had poor level of knowledge compared with respondents from Addis Ababa (Finfinnee), the capital (Table 4).

**Table 4.** Logistic regression analysis of odds ratio for knowledge of the respondents in relation to COVID-19 potential risk factors.

| 4. Demographic variables |  | Univariable logistic regression |  |  | Multivariable logistic regression |  |  |
| --- | --- | --- | --- | --- | --- | --- | --- |
|  |  | Crude OR | 95% CI | P-value | Adjusted OR | 95% CI | P-value |
| Sex | Male | 1.000 |  |  | 1 |  |  |
|  | Female | 0.921 | 0.747-1.135 | 0.440 | 0.864 | 0.7234-1.0923 | 0.621 |
| Age in ears | 18-29 | 1.000 |  |  | 1 |  |  |
|  | 30-39 | 0.829 | 0.637-1.080 | 0.165 | 0.923 | 0.6754-1.1073 | 0.243 |
|  | 40-49 | 0.944 | 0.675-1.320 | 0.735 | 1.021 | 0.6891-1.4225 | 0.14 |
|  | 50-59 | 0.473 | 0.307-0.728 | 0.001* | 0.562 | 0.4285-0.708 | 0.004* |
|  | 60+ | 0.290 | 0.154-0.545 | 0.000* | 0.387 | 0.1924-0.4982 | 0.001* |
| Residence | Urban | 1.000 |  |  |  |  |  |
|  | Rural | 0.682 | 0.541-0.860 | 0.001* | 0.772 | 0.5640-0.9245 | 0.004* |
|  | Single | 1.000 |  |  |  |  |  |
| Marital status | Married | 0.524 | 0.425-0.646 | 0.000* | 1.023 | 0.8776-1.2113 | 0.061 |
|  | Widow | 0.000 | 0.621-1.135 | 0.998 | 0.8823 | 0.7645-0.9921 | 0.12 |
|  | Farmer | 1.000 |  |  |  |  |  |
| Occupation | Government Employee | 20.599 | 12.681-33.461 | 0.000* | 16.4321 | 11.4564-27.9687 | 0.001* |
|  | Self-employed | 2.206 | 1.376-3.536 | 0.001* | 1.76 | 1.0248-2.6843 | 0.008* |
|  | Student | 8.716 | 5.512-13.784 | 0.000* | 5.0264 | 3.0245-10.2318 | 0.001* |
| Education | Illiterate | 1.000 |  |  |  |  |  |
|  | Basic | 2.023 | 1.181-3.466 | 0.000* | 1.276 | 0.9876-2.1346 | 0.086 |
|  | Primary | 4.327 | 2.447-7.654 | 0.000* | 3.0341 | 1.9842-5.6742 | 0.02* |
|  | Secondary | 3.740 | 2.197-6.366 | 0.000* | 3.042 | 2.4420-6.2143 | 0.012* |
|  | college/university | 16.434 | 10.335-26.130 | 0.000* | 14.026 | 9.56431-21.8764 | 0.001* |
|  | Addis Ababa | 1.000 |  |  |  |  |  |
| Region/chartered city | Afar | 0.745 | 0.569-1.102 | 0.091 | 0.703 | 0.5423-1.097 | 0.110 |
|  | Amhara | 0.513 | 0.349-0.756 | 0.001* | 0.497 | 0.2990-0.6542 | 0.007* |
|  | Benishangual Gumuz | 0.596 | 0.302-1.259 | 0.127 | 0.586 | 0.2882-0.7549 | 0.340 |
|  | Dire Dawa | 0.999 | 0.552-1.807 | 0.997 | 0.992 | 0.5286-1.577 | 0.481 |
|  | Harari | 0.696 | 0.306-1.159 | 0.427 | 0.621 | 0.2764-1.112 | 0.67 |
|  | Gambela | 0.743 | 0.431-1.280 | 0.284 | 0.709 | 0.476-1.1187 | 0.41 |
|  | Oromia | 0.470 | 0.305-0.889 | 0.000* | 0.4512 | 0.3218-0.8026 | 0.001* |
|  | Somali | 0.908 | 0.518-1.592 | 0.737 | 0.9 | 0.6234-1.4410 | 0.820 |
|  | SNNPRS | 0.553 | 0.334-0.918 | 0.022* | 0.54 | 0.3491-0.8823 | 0.230 |
|  | Tigray | 1.672 | 1.061-2.634 | 0.027 | 1.02 | 0.7465-3.2438 | 0.160 |

OD= odds ratio; CI= confidence interval; \* significant at $P \leq 0.05$

### 3.3. Attitude

Five questions were used to measure the attitude of the respondents towards COVID-19. More than 50% of the respondents either agree or strongly agree that traditional herbs and religious faith such as holy water can cure COVID-19. Half of the respondents think that it is unlikely to get sick from COVID-19. About 47% of the respondents do not agree with the principles of COVID-19 prevention methods such as hand washing, social distancing, avoiding non-essential travel and self-isolation (if have symptoms) and 57.1% of them disagree with the current introduction of state emergency to curb the spread of COVID-19. Only 40.4% of the respondents are confident that the government can prevent a nationwide outbreak of the disease (Table 5).

**Table 5.** Summary of respondents’ response to COVID-19 attitude questions (N=1570).

| no | Attitude questions | Response n (%); N= 1570 |  |  |  |
| --- | --- | --- | --- | --- | --- |
|  |  | Strongly disagree | Disagree | Agree | Strongly agree |
| 1 | I believe that traditional herbs and religious faith such as Holy water can cure COVID-19 | 588 (37.5) | 186 (11.8) | 290 (18.5) | 506 (32.2) |
| 2 | I am in favour of state of emergency introduced to control the spread of the coronavirus | 678 (43.2) | 220 (14.0) | 328 (20.9) | 344 (21.9) |
| 3 | I don't think that I will get sick from the coronavirus | 490 (31.2) | 322(20.5) | 286(18.2) | 472 (30.1) |
| 4 | I am confident that the government can prevent a nationwide outbreak of the coronavirus | 598 (38.1) | 388 (21.5) | 232 (14.8) | 402 (25.6) |
| 5 | Hand washing, social distancing, avoiding non-essential travel and self-isolation (if have symptoms) greatly help to contain the spreading of coronavirus | 386 (24.6) | 356 (22.7) | 262 (16.7) | 566 (36.1) |

The association between the socio-demographic variables and COVID-19 attitude was assessed by chi-square test (Table 5). The overall favourable attitude response rate was 43.8%. Respondents in the age group of 40-49 years had more favourable attitude (50.6%) towards COVID-19 prevention methods and principles than the other age groups (*χ*^2^=8.6, P ≤ 0.05). Interestingly rural residents had more favourable attitude (59.2%) than the corresponding urban residents (37.1%) (*χ*^2^ =65.996, P ≤ 0.001). There was strong association between occupation and attitude towards COVID-19 (*χ*^2^ = 335.735, P ≤ 0.0001). While government employees had the highest favourable response (77.8%), self-employed respondents had the least favourable response rate (15.7%). The educational status of the respondents was also found to be strongly associated with the attitude of the respondents towards COVID-19 (*χ*^2^=195.914, P ≤ 0.001). Respondents from Tigray (65.3%) and Amhara (63.2%) regions had relatively higher favourable attitude while respondents from Gambela (17.6%) and Afar (28.6%) had the lowest favourable attitude (*χ*^2^ =89.64, P≤0.001) (Table 6).

**Table 6.**
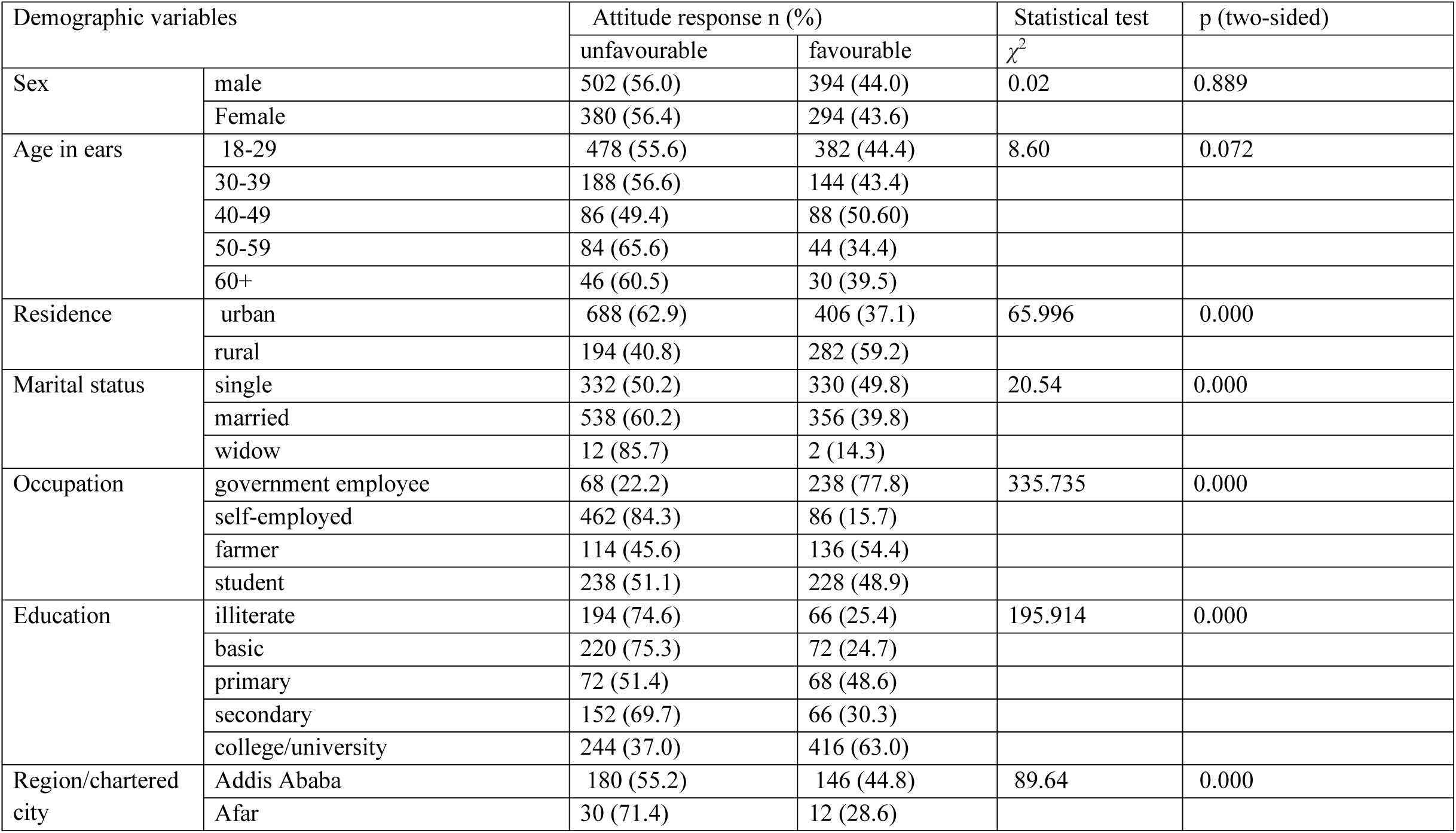
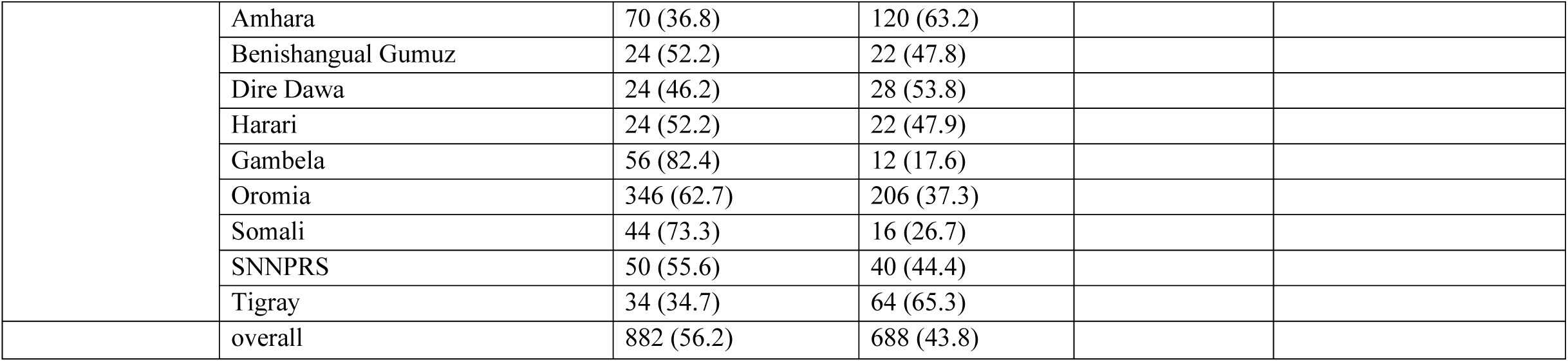
Respondents attitude score of COVID-19 by demographic variables (N-=1570).

Multivariable regression analysis has shown that residence, marital status, education, occupation and age are factors that most predict the attitude of the respondents with the correct predication of 73.2%. The odds of favourable attitude for being rural resident is twice as high as being urban resident (AOR=1.909; 95%CI: 1.436-2.538). On the other hand, being married (AOR=0.58; 95%CI: 0.398-0.854) and widow (AOR= 0.17; 95% CI: 0.034-0.854) were less likely to have favourable attitude compared to unmarried (single) respondents. Government employees had the highest odds of favourable attitude (OR=4.8) compared to farmers. As the educational level of the respondents increases, the odds of favourable attitude increase (OR=1.41, 1.68 and 3.01) for primary, secondary and college/university education respectively). While, Odds of favourable attitude for respondents from Amhara region is twice the odds of favourable attitude of respondents from Addis Ababa. The odds of favourable attitude of respondents from Afar, Gambela, Somali and Oromia are significantly low (P≤0.05) (Table 7).

**Table 7.** Logistic regression analysis of odds ratio for attitude of the respondents in relation to COVID-19 potential risk factors.

| Demographic variables |  | Favourable attitude towards Covid-19 prevention measures |  |  |  |  |  |
| --- | --- | --- | --- | --- | --- | --- | --- |
|  |  | Univariable binary logistic regression |  |  | multivariable logistic regression |  |  |
|  |  | Crude OR | 95% CI | P-value | adjusted OR | 95% CI | P-value |
| Sex | male | 1.00 |  |  |  |  |  |
|  | Female | 1.04 | 0.83-1.24 | 0.889 | 0.908 | 0.712-1.157 | 0.436 |
| Age in ears | 18-29 | 1.00 |  |  |  |  |  |
|  | 30-39 | 0.96 | 0.958-1.237 | 0.745 | 0.87 | 0.623-1.234 | 0.531 |
|  | 40-49 | 1.28 | 0.924-1.774 | 0.138 | 1.03 | 0.7834-2.292 | 0.643 |
|  | 50-59 | 0.66 | 0.444-0.967 | 0.033* | 0.72 | 0.548-1.009 | 0.090 |
|  | 60+ | 0.82 | 0.505-1.318 | 0.406 | 0.89 | 0.728-1.233 | 0.620 |
| Residence | Urban | 1.00 |  |  |  |  |  |
|  | Rural | 2.46 | 1.977-3.070 | 0.000* | 1.909 | 1.436-2.538 | 0.000* |
| Marital status | Single | 1.00 |  |  |  |  |  |
|  | Married | 0.61 | 0.435-0.855 | 0.00* | 0.583 | 0.398-0.854 | 0.000* |
|  | Widow | 0.39 | 0.078-1.954 | 0.25 | 0.172 | 0.034-0.854 | 0.006* |
| Occupation | Farmer | 1.00 |  |  |  |  |  |
|  | Government employee | 2.79 | 1.926-4.031 | 0.000* | 4.758 | 2.144-7.321 | 0.000* |
|  | Self employed | 0.15 | 0.106-0.211 | 0.000* | 0.559 | 0.423-0.992 | 0.042* |
|  | Student | 0.53 | 0.351-0.808 | 0.003* | 0.852 | 0.582-1.113 | 0.110 |
| Education | Illiterate | 1.00 |  |  |  |  |  |
|  | Basic | 0.96 | 0.654-1.415 | 0.844 | 0.88 | 0.6783-1.194 | 0.740 |
|  | Primary | 2.78 | 1.800-4.282 | 0.000* | 1.41 | 1.09-3.247 | 0.023* |
|  | Secondary | 1.28 | 0.854-1.908 | 0.234 | 1.68 | 1.420-2.228 | 0.040* |
|  | College/university | 5.01 | 3.636-6.908 | 0.000* | 3.01 | 2.078-5.193 | 0.000* |
| **Region/chartered city** | Addis Ababa | 1.00 |  |  | 1 |  |  |
|  | Afar | 0.49 | 0.244-0.997 | 0.049* | 0.52 | 0.242-0.8812 | 0.024* |
|  | Amhara | 2.11 | 1.465-3.050 | 0.000* | 1.876 | 1.0429-3.378 | 0.036* |
|  | Benishangual Gumuz | 1.13 | 0.609-2.097 | 0.698 | 0.597 | 0.267-1.332 | 0.208 |
|  | Dire Dawa | 1.44 | 0.799-2.588 | 0.225 | 0.841 | 0.499-1.416 | 0.514 |
|  | Harari | 1.14 | 0.719-2.897 | 0.698 | 0.820 | 0.371-1.812 | 0.624 |
|  | Gambela | 0.26 | 0.136-0.0511 | 0.000* | 0.259 | 0.113-0.595 | 0.001* |
|  | Oromia | 0.73 | 0.556-0.969 | 0.029* | 0.537 | 0.325-0.888 | 0.015* |
|  | Somali | 0.45 | 0.243-0.827 | 0.010* | 0.245 | 0.112-0.535 | 0.000* |
|  | SNNPRS | 0.99 | 0.617-1.577 | 0.954 | 0.702 | 0.368-1.374 | 0.302 |
|  | Tigray | 2.32 | 1.451-3.712 | 0.000* | 1.016 | 0.473-2.181 | 0.968 |

### 3.4. Practice

The practice of the respondents to prevent COVI-19 was assessed by 5 questions. About 43% of the respondents never practice any of the COVID-19 prevention methods and only less than one fifth (19.1%) of the respondents follow COVID-19 prevention measures either usually or always. About a third (31.0%) of respondents never wash their hands with soap and 42.9% of the respondents never avoided touching their eyes, nose, and mouth with their unwashed or unsensitized hand. Four of five respondents (81.4%) do not keep 2-meter social distancing whenever they are out for essential reasons. More than half of the respondents never avoided social gatherings for religious, festivals, or other ceremonies since the introduction of the state of emergency (Table 8).

**Table 8.** Summary of respondents’ response to COVID-19 practice questions (N=1570).

| no | Practice questions | response n (%); N= 1570 |  |  |  |
| --- | --- | --- | --- | --- | --- |
|  |  | never | sometimes | usually | always |
| 1 | I wash my hands with soap frequently | 486 (31.0) | 630 (40.1) | 344 (21.9) | 110 (7.0) |
| 2 | I avoided non-essential travel since the lockdown | 682 (43.4) | 598 (38.1) | 214 (13.6) | 76 (4.8) |
| 3 | I always keep the social distancing of at least 2-meters whenever I am out for essential reasons | 710 (45.2) | 568 (36.2) | 228 (14.5) | 64 (4.1) |
| 4 | I avoided social gatherings for religious, festivals, or other ceremonies since the state emergency is introduced | 810 (51.6) | 508(32.4) | 190 (12.1) | 62 (3.9) |
| 5 | I avoid touching my eyes, nose, and mouth with my unwashed hands. | 678 (43.2) | 686 (43.7) | 160 (10.2) | 46 (2,9) |
|  | Overall | 673 (42.9) | 598 (38.1) | 227 (14.5) | 72 (4.6) |

There was no statistically significant difference between gender, and among age and marital status in the implementation of COVID-19 preventive measures at individual level. On the other hand, there is statistically significant difference in practicing COVID-19 preventive measure among residence, education, occupation and region of the respondents (P≤0.001). While 27.6% of urban respondents practice COVID-19 prevention measures, only 16.8% of the corresponding rural residents practice these measures either usually or always. Only a small proportion of respondents practice COVID-19 prevention measures in SNNP, Gambela and Dire Dawa compared to respondents from other regions (Table 9).

**Table 9.** Respondents practice score of COVID-19 by demographic variables (N-=1570).

| Demographic variables |  | Response n (%) |  | Statistical test | P (two-sided) |
| --- | --- | --- | --- | --- | --- |
| | | Never/sometimes | Usually/always | $\chi^2$ | |
| Sex | Male | 663 (74.0) | 233(26.0) | 3.174 | 0.085 |
|  | Female | 525 (77.9) | 149(22.1) |  |  |
| Age in ears | 18-29 | 652 (75.8) | 208 (24.2) | 0.452 | 0.978 |
|  | 30-39 | 249 (75.0) | 83 (25.0) |  |  |
|  | 40-49 | 132 (75.9) | 42 (24.1) |  |  |
|  | 50-59 | 99 (77.3) | 29 (22.7) |  |  |
|  | 60+ | 56 (73.7) | 20 (26.3) |  |  |
| Residence |  |  |  |  |  |
|  | Urban | 792 (72.4) | **302 (27.6)** | **21.007** | **0.000** |
|  | Rural | 396 (83.2) | 80 (16.8) |  |  |
| Marital status | Single | 486 (73.4) | 176 (26.6) | 3.429 | 0.180 |
|  | Married | 692 (77.4) | 202 (22.6) |  |  |
|  | Widow | 10 (71.4) | 4 (28.6) |  |  |
| Occupation | Government employee | 186 (60.8) | 120 (39.2) | 57.476 | 0. 000 |
|  | Self-employed | 423 (77.2) | 125 (22.8) |  |  |
|  | Farmer | 219 (87.6) | 31 (12.4) |  |  |
|  | Student | 360 (77.3) | 106 (22.7) |  |  |
| Education | Illiterate | 213 (81.9) | 17 (18.1) | 3.056 | 0.000 |
|  | Basic | 240 (82.2) | 52 (17.8) |  |  |
|  | Primary | 114 (81.4) | 26 (18.6) |  |  |
|  | Secondary | 166 (76.1) | 52 (23.9) |  |  |
|  | College/university | 455 (68.9) | 205 (31.1) |  |  |
| **Region/chartered city** | Addis Ababa | 185 (56.7) | 141 (43.3) | 95.236 | 0.000 |
|  | Afar | 36 (85.7) | 6 (14.3) |  |  |
|  | Amhara | 155 (81.6) | 35 (18.4) |  |  |
|  | Benishangual Gumuz | 36 (78.3) | 10 (21.7) |  |  |
|  | Dire Dawa | 46 (88.5) | 6 (11.5) |  |  |
|  | Harari | 39 (84.8) | 7 (15.2) |  |  |
|  | Gambela | 60 (88.2) | 8 (11.8) |  |  |
|  | Oromia | 434 (78.6) | 118 (21.4) |  |  |
|  | Somali | 48 (80.0) | 2 (20.0) |  |  |
|  | SNNPRS | 80 (88.9) | 10 (11.1) |  |  |
|  | Tigray | 69 (70.4) | 29 (29.6) |  |  |
|  | Overall | 1188 (75.7) | 382 (24.3) |  |  |

Multivariable logistic regression analysis has shown that residence, region, education and occupation are the most predicting factors (76.1%) of COVID-19 appropriate practical measures. Being rural resident has a probability of 43.0% not to practice COVID-19 prevention measures than being urban resident. The odds of government employee are more than twice (AOD=2.268) compared with the odds of being farmer towards appropriate COVID-19 practical measures while the odds of being self-employed was not statistically significantly different compared with the odds of being a farmer. The odds of respondents from all other regions were significantly lower than the odds of Addis Ababa (Finfinnee) respondents (Table 10).

**Table 10.** Logistic regression analysis of odds ratio for practice of the respondents in relation to COVID-19 potential risk factors.

| Demographic variables |  | Good practice towards Covid-19 prevention |  |  |  |  |  |
| --- | --- | --- | --- | --- | --- | --- | --- |
|  |  | Univariable binary logistic regression |  |  | multivariable logistic regression |  |  |
|  |  | Crude OR | 95% CI | P-value | adjusted OR | 95% CI | P-value |
| Sex | male | 1 |  |  | 1 |  |  |
|  | Female | 0.808 | 0.638-1.022 | 0.075 | 0.852 | 0.661-1.099 | 0.218 |
| Age in ears | 18-29 | 1 |  |  | 1 |  |  |
|  | 30-39 | 1.045 | 0.779-1.401 | 0.769 | 1.860 | 1.003-3.447 | 0.049 |
|  | 40-49 | 0.997 | 0.682-1.459 | 0.989 | 1.587 | 0.861-2.995 | 0.139 |
|  | 50-59 | 0.918 | 0.590-1.429 | 0.705 | 1.643 | 0.858-3.146 | 0.134 |
|  | 60+ | 1.120 | 0.656-1.909 | 0.679 | 1.344 | 0.683-2.645 | 0.391 |
| Residence | urban | 1 |  |  | 1 |  |  |
|  | rural | 0.530 | 0.403-0.697 | 0.000* | 0.570 | 0.413-0.786 | 0.001* |
| Marital status | single | 1 |  |  | 1 |  |  |
|  | married | .806 | 0.639-1.018 | .070 | 1.224 | 0.342-4.383 | 0.757 |
|  | widow | 1.105 | 0.342-3.567 | .868 | 1.700 | 0.496-5.821 | 0.398 |
| Occupation | farmer | 1 |  |  |  |  |  |
|  | government employee | 4.558 | 2.934-7.081 | 0.000* | 2.268 | 1.519-3.385 | 0.000* |
|  | Self-employed | 2.088 | 1.364-3.196 | 0.001* | 1.102 | 0.584-2.081 | 0.765 |
|  | student | 2.080 | 1.348-3.211 | 0.001* | 1.372 | 0.875-2.151 | 0.168 |
| Education | illiterate | 1.000 |  |  |  |  |  |
|  | basic | 0.982 | 0.635-1.518 | 0.935 | 2.093 | 1.3923-3.146 | 0.000* |
|  | primary | 1.034 | 0.608-1.757 | 0.903 | 2.283 | 1.561-3.338 | 0.000* |
|  | secondary | 1.420 | 0.911-2.212 | 0.122 | 2.343 | 1.461-3.758 | 0.000* |
|  | college/university | 2.042 | 1,430-2.916 | 0.000* | 1.494 | 1.036-2.153 | 0.031* |
|  | Addis Ababa | 1 |  |  | 1 |  |  |
| Region/chartered city | Afar | 0.2187 | 0.090-0.533 | 0.001* | 0.559 | 0.337-0.928 | 0.025* |
|  | Amhara | 0.2963 | 0.193-0.454 | 0.000* | 0.236 | 0.095-0.585 | 0.002* |
|  | Benishangual Gumuz | 0.364 | 0.175-0.759 | 0.007* | 0.367 | 0.234-0.573 | 0.000* |
|  | Dire Dawa | 0.1710 | 0.071-0.412 | 0.000* | 0.519 | 0.371-0.727 | 0.000* |
|  | Harari | 0.235 | 0.102-0.452 | 0.001* | 0.250 | 0.108-0.582 | 0.001* |
|  | Gambela | 0.175 | 0.081-0.378 | 0.000* | 0.211 | 0.103-0.431 | 0.000* |
|  | Oromia | 0.357 | 0.265-0.481 | 0.000* | 0.395 | 0.199-0.785 | 0.008* |
|  | Somali | 0.328 | 0.168-0.641 | 0.001* | 0.234 | 0.107-0.515 | 0.000* |
|  | SNNPRS | 0.164 | 0.082-0.328 | 0.000* | 0.406 | 0.191-0.861 | 0.019* |
|  | Tigray | 0.551 | 0.339-0.896 | 0.016* | 0.154 | 0.064-0.373 | 0.000* |

## 4. Discussions

COVID-19 is an infectious disease caused by sever acute respiratory coronavirus 2 (SARS-CoV-2) and its being pandemic poses a significant threat to public health (9). Given that the disease is pandemic threat with no vaccine or proven treatment drug, preventive measures are the most essential methods available to reduce the infection rates and control the spread of the disease. This implies that public adherence to preventive and control measures are essential to curb the disease. The extent of public adherence to preventive and control measures is affected by their knowledge, attitude and practice (KAP). Therefore, this study was envisioned to assess the KAP of the Ethiopian population for the control of COVID-19.

The finding has shown that the participants level of COVID-19 knowledge is unsatisfactory. Only 42% of the respondents achieved a satisfactory knowledge score and the result is lower than most previous reports from various countries (2, 13, 19, 23-25). Our study has shown that Ethiopians had lower level of COVID-19 knowledge than Chinese (80.5%) ((25), Saudi Arabia (81.6%)(24), Bangladesh (54·87%) (26), Malaysia (80.5%) (23) and Tanzania (84.4%) (27) but similar with a study reported from Cameroon (43,7%) (28).

Poor level of knowledge of Ethiopians towards COVID-19 could make the spread of the pandemic worst coupled with the country’s limited capacity of testing, contact tracing and isolation of suspected cases. Until the end of June 2020, Ethiopia has tested only less than 300,000 peoples among the 114 million population indicating that Ethiopia is far from achieving the pandemic control.

Knowledge scores significantly differ across socio-demographic factors which underlines the need for targeted public awareness campaign. Although most of the respondents (73.6%) have heard about COVID-19, only 42.4% and 37.8% knew COVID-19 transmission and clinical manifestations, respectively. Furthermore, only less than 20% of the respondents knew that currently there is no effective drug to cure COVID-19 nor vaccines to prevent the infection. This poor level of knowledge towards COVID-19 could be due to limited engagement of Ethiopian institutions, religious sectors, political parties and policy-decision makers to generate contextual knowledge platform to provide an easy and accessible ways of getting credible information. It could be also due to top-down community awareness campaigns, lack of engaging diverse groups by the government, communication overload as large and conflicting information are directed at federal, regional state, and local health bureau from multiple sources which challenges the credibility and acceptability of the information by the general public. However, these proposed scenarios need further investigation.

Multivariable logistic regression analysis has shown that respondents from Oromia and Amhara regions are 55% and 50% more likely to be less knowledgeable about COVID-19 respectively than other regions. The poor level of COVID-19 knowledge in these two regions could be due to the vastness of these regions to address every corner of the regions adequately using the limited resources and infrastructure the country has. The sporadic protests and instability in these two regions cannot also be ignored affecting the consolidation of the information delivered from the government. This finding is very important since it may indicate that more effort should be exerted to deliver the desired message in these two most populous regions of the country, which account for more than 65% of the population of the country, (29) to control the spread of the disease.

Majority of the respondents (59.6%) have no confidence on the government to control the nationwide outbreak of the disease and 57.1% of the respondents disagree with the current introduction of state of emergency to curb the spread of COVID-19 by the government. Unexpectedly, half of the respondents do not agree with the principles of COVID-19 prevention methods such as hand washing, social distancing, avoiding non-essential travel and self-isolation symptomatic patients. As a result, only one in five respondents practice COVID-19 prevention measures satisfactorily. It is very difficult to explain why the attitude and practice of the public to fight COVID-19 is low unlike in other countries such as China (25), Egypt (30), Sudan (31) and Tanzania (27). One reason could be the top-down flow of government procedures to contain the virus may be perceived as authoritarian attempts by government to consolidate control, leading to a loss of faith in the government. This pinpoints that the government need to re-examine the mode of delivery of COVID-19 related messages to the public. Another reason could be related to poverty and food insecurity. The state of emergency weigh most heavily on the poor, who are often part of the informal economy and thus dependent on day-to-day making money to feed themselves and their families. For many of them, a day without work means a day without food. Hence, they may resist to follow government advice.

The attitude of the respondents toward COVID-19 significantly varies across some sociodemographic factors. Although rural residents had lower level of COVID-19 knowledge, a significantly higher proportion of (59.2%) them had favourable attitude than urban residents (37.1%) towards COVID-19. Multivariate logistic regression analysis has shown that the odds of favourable attitude of being a rural resident was twice than being an urban resident. This could be due to the fact that rural residents have limited access to various information sources particularly on social media (32). Hence, they largely depend on the information from the local and national governors. Rural residents are also less likely than the urban residents to translate service dissatisfaction into discontent with their government and hence have more trust in government and evaluate information from local and national officials more positively than urban peers (33).

Only one in 5 of the respondents adhered to COVID-19 prevention measures either usually or always. While a third of respondents never wash their hands with soap, more than 80% of the respondents had never applied a 2-meter social distancing rule at least once. Ethiopia is among one of the poorest sub-Sharan countries where a significant proportion of the population (70-80%) have no access to adequate water supply, sanitation and hygiene facilities (34) which challenges the handwashing practice to limit the spreading of COVID-19. Maintaining social distance is unattainable practice in Ethiopia since majority of the citizens live in slums. About 80% of Addis Ababa, the capital of the country, is considered slum areas, characterized by widespread sanitation challenges where families live in crowded rooms and are exposed to health and safety risks (35).

The ability of the government to persuade people to internalize the externality they would impose on the community by not reducing their mobility is one of the most important elements of compliance (36). This is particularly important where the ability to enforce lockdowns on a large scale is unattainable. Evidence from different countries indicates that political beliefs, coupled with differences in media consumption, have important implications for risk perceptions and compliance with social distancing and the Ethiopian government should revisits its COVID-19 prevention strategy (36).

Strengthening of government measures, community-based sensitization and health education programs will go a long way to prepare the community to overcome the pandemic. Providing this education to the population will help fill the knowledge gaps reflected and correct the misperception regarding the attitude of the population towards the disease which ultimately improves the practice.

## 5. Conclusions and recommendations

In summary, the present study was able to provide a comprehensive investigation of the knowledge, attitude and practice of Ethiopians toward COVID-19. The finding suggest that Ethiopians have inadequate level of knowledge on COVID-19 and are generally have a mixed outlook on overcoming the pandemic. The poor adherence to COVID-19 prevention practice by the public underlines the need for urgent reinforcing preventive measures and intensifying sensitization campaigns to fill the knowledge gap of the population with regards to COVID-19 and to persuade people to follow the preventive measures set by the government. At the same time, there is a need to evaluate the impact of these measures on the knowledge and practices of the population with the progression of the pandemic in Ethiopia. Finally, we believe that the study will inspire the healthcare authorities, and media to spread more COVID-19 related accurate knowledge which ultimately results in better preventive practices toward COVID-19. Since no proven medicine or vaccine is developed yet, the best way to curb the spreading of the disease is maximizing knowledge and preventive practices toward COVID-19 at all levels.

